# A miniature cellulosome with novel interaction modes

**DOI:** 10.1101/2023.01.24.524588

**Authors:** John Allan, Gary W Black

## Abstract

Cellulosomes are efficient enzymatic nanomachines which have arisen for the degradation of cellulosic biomass. They are found abundantly in soil-dwelling microbes and bacteria which thrive in the stomachs of ruminant mammals. Two protein domains, cohesins and dockerins, characterise cellulosomes. These domains interact with each other to form enormous complexes with as many as 130 individual proteins. Annotation of the genome of *Cellulosilyticum lenotcellum* DSM 5427 revealed one single cohesin and one single dockerin domain. This pales in comparison to most cellulosomal organisms. We have recombinantly produced the proteins which bear these domains and demonstrated cellulase activity, which is enhanced by interaction between the two proteins. Moreover we have identified additional novel interacting partners for this unique cellulosome complex. This broadens the repertoire and definitions of cellulosomes overall.

## Introduction

*Cellulosilyticum lentocellum* was first isolated from a riverbank washed by effluent from a nearby paper mill in the River Don, in Aberdeenshire, Scotland^1^. Since its initial culture-based isolation however there have been a dearth of focussed studies on its physiology and molecular microbiology. It has been identified in various other environments, for example the faeces of pangolins^2^ and giant pandas^3^, sometimes enriched in landfill sites^4^ or activated sludge^5^. It has also been explored biotechnologically in astronautical hydroponics projects^6^, and some of its epimerases have been used in the production of prebiotics^7^.

Clearly, *C. lentocellum* is a biotechnologically useful organism for the handling of plant biomass and other lignocellulosic materials. Further understanding of this organism is required to realise its evident biolotechnological potential.

As a former *Clostridia, C. lentocellum* is closely related to many cellulosome utilising organisms^1,8,9^. Cellulosomes are complex enzymatic apparatuses secreted from cells and, in some cases, anchored to cell surfaces. Their function is cellulose and hemicellulose degradation to simple sugars, that can be utilised by bacterial cells^10^. They are comprised of enzymes which contain dockerin domains, and scaffoldin proteins which contain cohesin domains, and in some cases also dockerins. The near-covalent strength interaction between cohesins and dockerins^11,12^ is the defining feature of cellulosomes and facilitates the assembly of complexes containing as many as 130 enzymes^13^. Cohesin-dockerin interactions are also almost species specific, in that cohesins from one organism may not interact with the dockerin of another. Some instances exist where this paradigm is broken^14^, whether or not this occurs in nature however remains to be seen.

Typically, enzymes within cellulosomes are cellulases and xylanases, however other enzymes have been described too. The canonical cellulosome is produced by *Acetivibrio thermocellus* (previously *Clostridium thermocellum*)^15^. This contains a scaffoldin anchored to the cell surface. In the canonical cellulosome of *A. thermocellus*, this is a small protein called OlpB which binds to the S-layer of the cell^16^. Cellulases and xylanases secreted with dockerin domains may interact with these cohesin domains, as well as additional dockerin containing scaffoldin proteins (termed type ii scaffoldins), giving rise to more complex branched structures.

Within the *C. lentocellum* genome, there is a putative miniature cellulosome encoded. There is a locus containing two putative CDS referred hereafter as ClSca and ClGH. Within each of which there are annotatable cohesin (CISca) and dockerin (ClGH) domains (Figure 1). BLAST Conserved Domain^17,18^ searches and SMART^19,20^ annotations predicted the presence cohesin and dockerin domains on these proteins.

**Figure 1.**
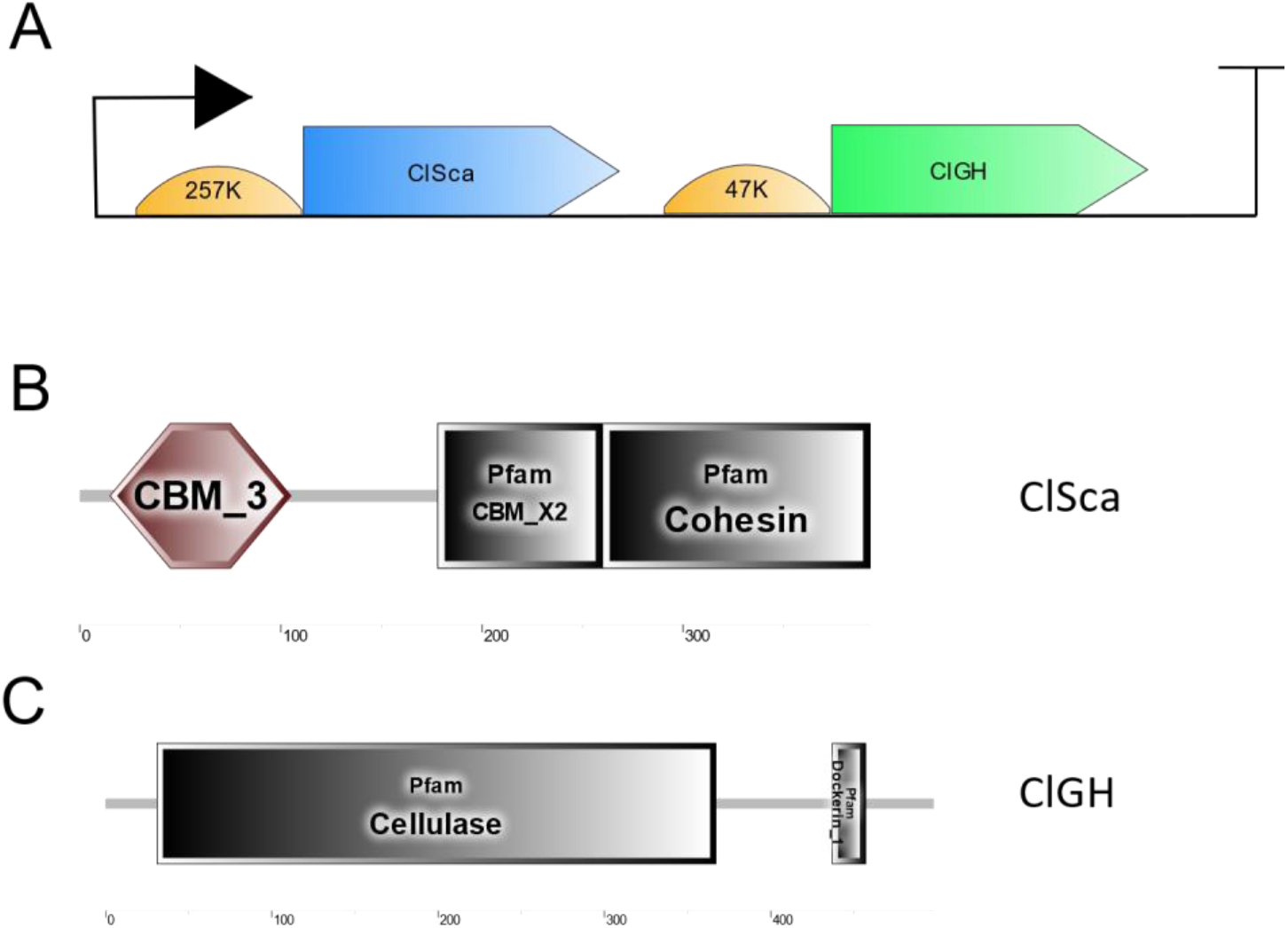
The *Cellulolosilyticum lentocellum* genome encodes an uncharacterised operon that contains novel cohesin and dockerin domains. By PSI-BLAST searching of genomes contained in the GEBA collection of microorganisms (DSMZ), a novel pair of cohesin and dockerin domains was identified. (A) Promoters and ribosome binding sites were annotated with the BPROM and RBS Calculator tools respectively. Putative domains were annotated using the BLAST conserved domain search, and EMBL SMART tools. SMART cartoon output is shown here for ClSca (B), and ClGH (C).

Phylogenetic analyses and sequence alignment of the sequences annotated in PFAM containing dockerin and cohesin domains shows that the *C. lentocellum* dockerin is most closely related to those of *Cellulosilyticum sp*. WCF-2 (Supplementary Figure 1). Alignments with characterised cohesin and dockerin domains (determined for this purpose as those with structures in the Protein Data Bank) indicate that the *C. lentocellum* cohesin domain is unique to others which have been described previously, while the dockerin is ∼60% similar to the A. *thermocellus* sequences which exists in the original, defining co-crystal structure of a cohesin-dockerin interaction. CISsa and CIGH also share close similarity with other indicated proteins (Supplementary Figure 1).

We set out here to define this putative cellulosome biochemically and explore its interactions with other elements of cellular physiology.

## Results

### Biochemical Characterisation of Putative Cellulosomal Proteins

In order to characterise this putatative cellulosomal operon, we amplified the genes using genomic DNA prepared from *C. lentocellum* strain DSM5427. These were ligated by classical methods into plasmid vectors which added translationally fused hexahistidine tags. The *clsca* gene was cloned into pQE80L to generate pQEClSca and CISca was produced at 500 mL scale by *E. coli* BL21(DE3) in autoinduction medium. The *cigh* gene was cloned into pET28a to generate pETClGHdsec and CIGH was produced at 50 mL scale by *E. coli* BL21(DE3) in autoinduction medium. Both were subsequently purified using TALON resin.

ClSca and ClGH each contain putative domain structures that are characteristic of cellulose proteins. ClGH contains a glycosyl hydrolase domain of class 3 (GH3), along with a dockerin domain. A putative carbohydrate binding module of class 3 (CBM3) is present on the ClSca polypeptide (Figure 1). This is a common feature of cellulosomal scaffoldin proteins. The domain permits a close interaction of the cellulosome complex with cellulose substrate chains. In order to first test the hypothesis that this is reflected in ClSca, the purified protein was incubated with insoluble, microcrystalline cellulose (Avicel PH-101, Sigma). Concentration gradients were considered for both protein and Avicel. After three hours, the Avicel was removed from the suspension by centrifugation, and washed with CDI buffer (see Experimental Procedures). Samples were prepared from the resulting supernatants and the Avicel pellet for SDS-PAGE and run on 15% polyacrylamide gels (Figure 2). ClSca was clearly seen to increase in abundance in the bound fractions as the concentration of Avicel was increased, and it was removed from the suspension by the Avicel regardless of its concentration.

**Figure 2.**
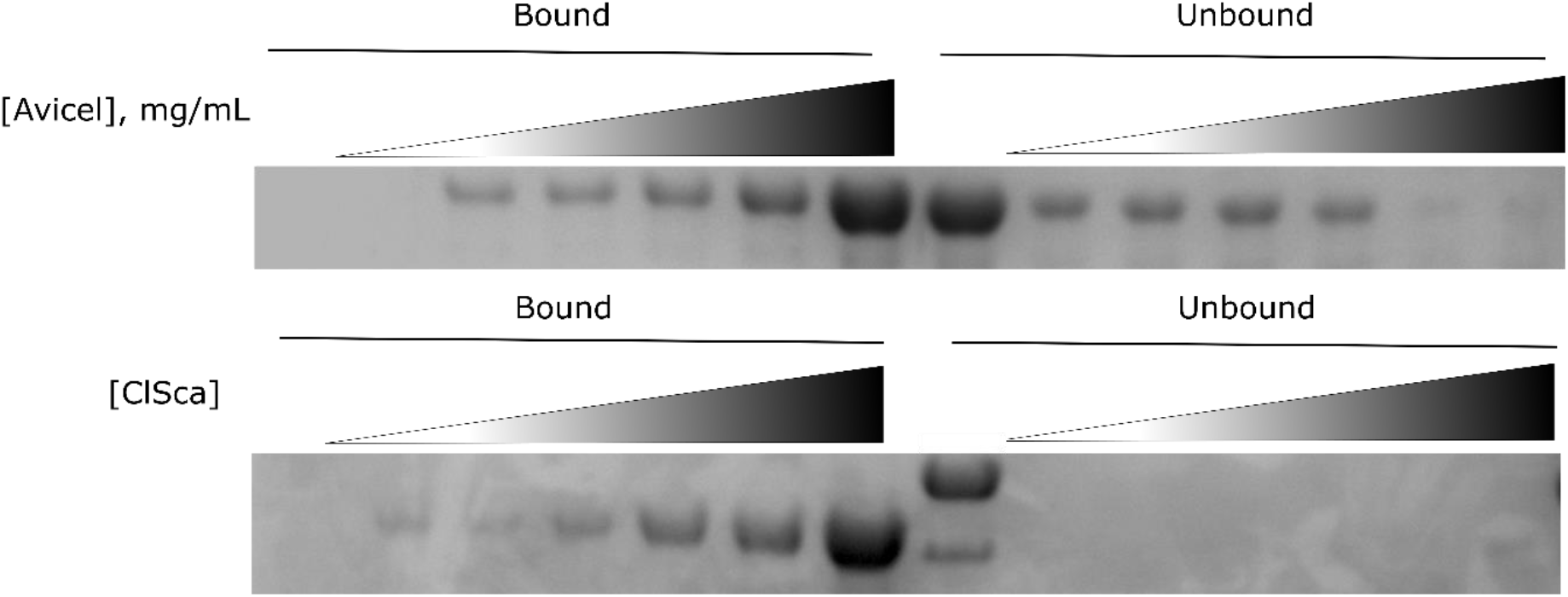
ClSca strongly interacts with microcrystalline cellulose. (A) Purified ClSca was purified and incubated with Av icel at concentrations between 0 and 50 mg/mL, with ClSca at 1.36 mg/mL in each replicate for three hours with rotary shaking at 4 oC. Samples were then centrifuged to separate Avicel from the soluble fraction. The soluble fraction was retained as unbound protein, and the insoluble Avicel fraction mixed with Laemmli buffer for SDS-PAGE analysis. (B) ClSca was incubated for three hours at 4 oC, while Avicel was retained at 25 mg/mL, ClSca was used in a range of concentrations between 1.3 mg/L and 0.13 mg/mL. Bound and unbound fractions were then obtained as in A.

Next, the activity of the dockerin-bearing glycosyl hydrolase, CIGH, was characterised. The purified protein was incubated for three hours with phosphoric acid swollen cellulose (PASC) at various concentrations and released reducing sugars measured by DNSA to determine K_m_ and Vmax constants of the enzyme. This revealed an incredibly high KM, of around 175 mg/mL suggesting a poor affinity of the enzyme for PASC (Figure 3A). Additionally, we probed the specificity of the binding pocket of the enzymes active site by testing the activity of the enzyme on glucose and relevant oligomers cellobiose, cellotriose, cellotetraose and cellohexaose. The oligosachharides released were analysed by HPAEC, shown in Figure 3B.

**Figure 3.**
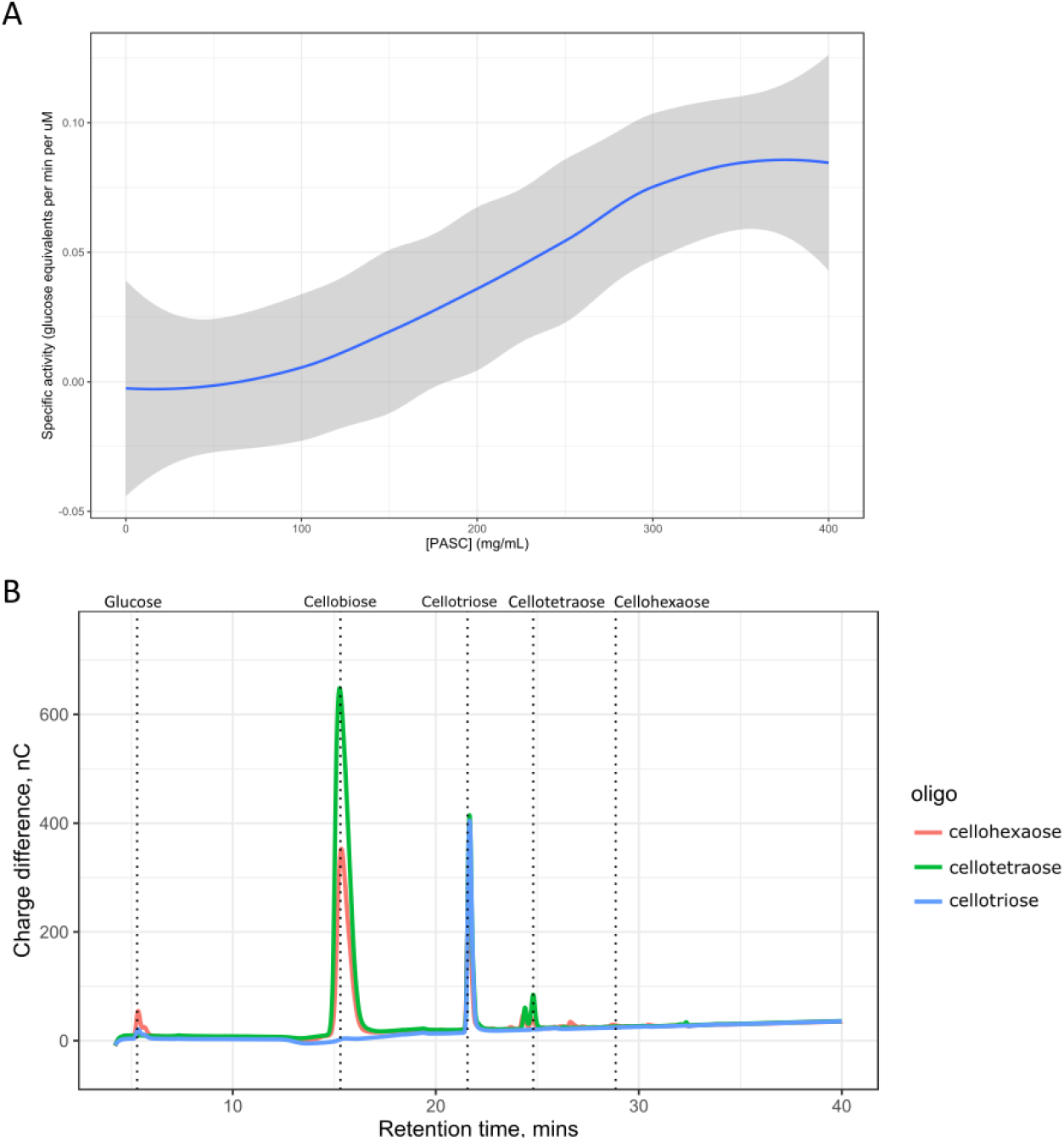
ClGH5 is a cellobiohydrolase capable of cellulose degradation. (A) ClGH5 was incubated with PASC for 2 hours at concentrations between 0 and 400 mg/mL and reducing sugars released measured by DNSA. This shows a K_M_ of around 175 mg//mL, and V_max_ of around 350 mg/mL. (B) Enzymatic assays were performed using cellobiose, cellotriose, cellotetraose and cellohexaose. The resulting sample was analysed by HPAEC, with representative traces plotted here, n = 3. Dotted lines mark the elution times of indicated standards.

The defining feature of the cellulosome is the interaction of the cohesin and dockerin domains. To test the hypothesis that the cohesin and dockerin domains present on ClSca and ClGH, respectively, can interact to form a miniature cellulosomal complex, the purified proteins were incubated at room temperature before being subjected to size-exclusion chromatography. The individual proteins were also analysed identically. When incubated together, a 90 kDa analyte was eluted, along with a smaller one. Individually, ClSca eluted at around 40 kDa and ClGH at around 55 kDa (Figure 4). Cellulase activity was retained in the 90 kDa fractions, as seen by TLC. Samples relating to each eluted protein were analysed via SDS-PAGE, revealing the 90 kDa fraction to be composed of a mixture of ClSca and ClGH. The smaller peak, which also retained cellulase activity, was slightly smaller than ClGH, and bigger than ClSca.

**Figure 4.**
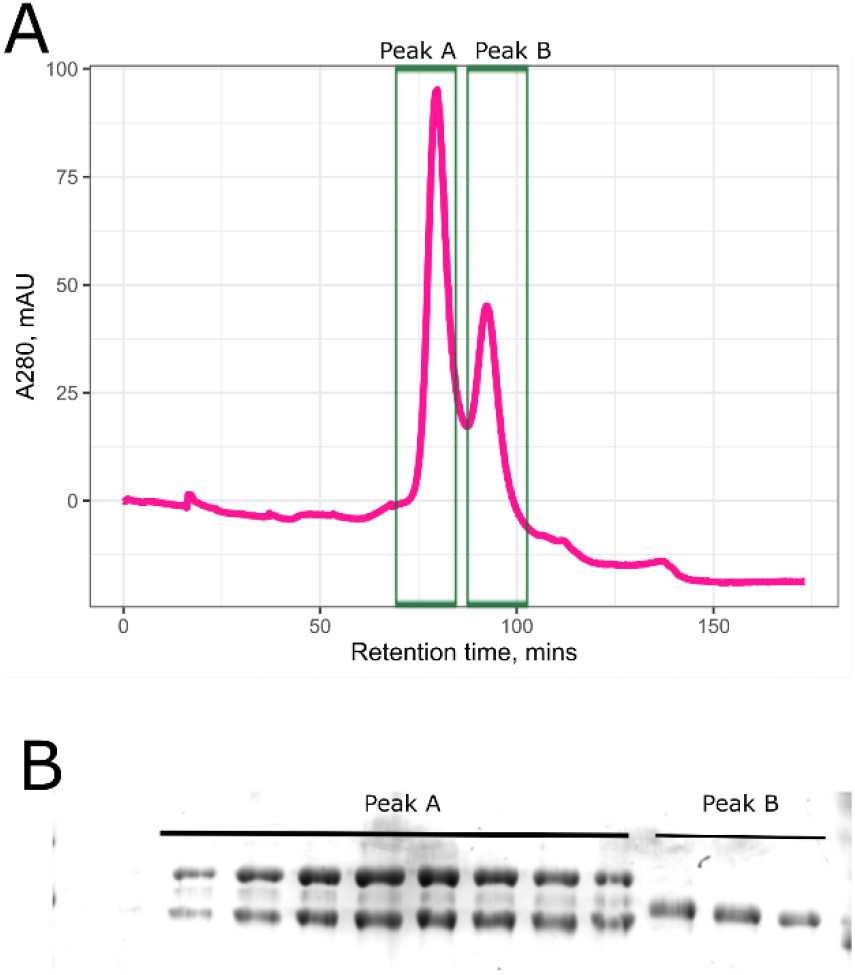
ClSca interacts with ClGH5. (A) ClSca and ClGH5 were purified in parallel and then concentrated ∼100-fold with a 30,000 MWCO centrifugal filter. Proteins were mixed at an equimolar ratio at 0.34 uM, and then applied to a calibrated 16/600 200 pg superdex size exclusion column. Samples collected at seleced peaks were also analysed by SDS-PAGE. (B) Samples were analysed by SDS-PAGE. Enzyme activity was also probed and assessed by TLC using these samples indicating ClGH activity in each fraction represented here, data not shown.

Satisfied that ClGH and ClSca form a complex, we next sought to understand how ClSca may affect the enzymatic activity of ClGH. The purified proteins were incubated at an equimolar ratio with either soluble carboxymethyl cellulose (CMC) or insoluble Avicel, at equal concentrations by mass. The activity was measured by DNSA, which indicated activity is much greater on the soluble CMC substrate, and in each case, it is enhanced by the addition of ClSca. However, the degree of enhancement was greater on the insoluble Avicel (∼3-fold) than on CMC (∼1.25-fold) (Figure 5).

**Figure 5.**
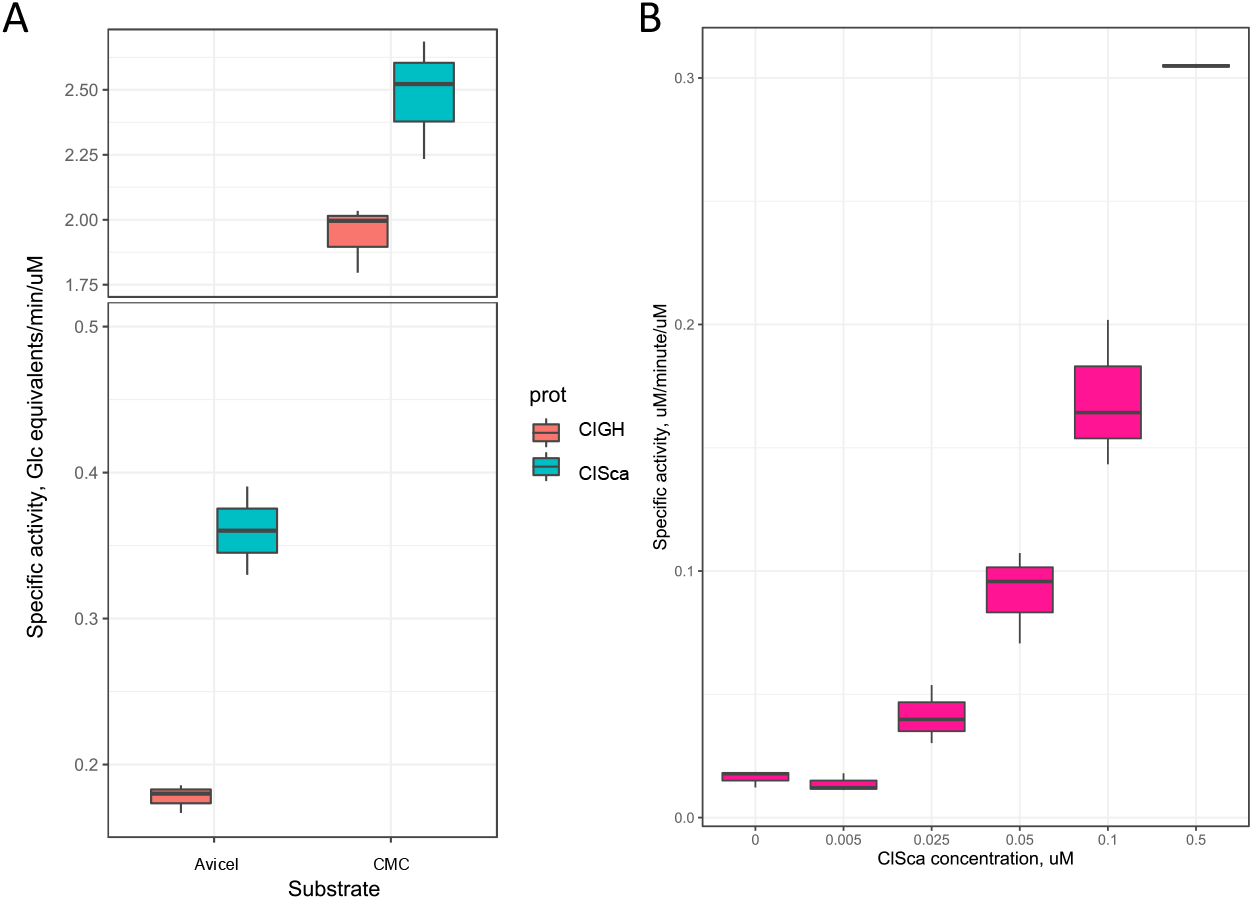
ClGH activity is enhanced upon addition of ClSca. Purified ClGH was incubated with ClSca at ratios from a 10-times excess of ClGH, to ten-times excess of ClSca, with ClGH retained at a concentration of 0.05 uM throughout. Released reducing sugars were assayed by DNSA as a measurement of total activity, while the results were in parallel validated with qualitative HPAEC.

### Co-precipitation of additional cellulosomal interacting partners

Since it was unusual that there is only one annotatable dockerin-bearing enzyme in the genome of *C. lentocellum* DSM5427, we hypothesised that there are additional interacting partners of ClSca. To test this hypothesis we performed a baited pull down experiment with purified ClSca as bait. Cell-free extract obtained from *E. coli* BL21(DE3) expressing ClSca was run through TALON resin, and washed with 10 mM imidazole in CDI buffer. Clarified cell-free extract of *C. lentocellum* grown with the addition of cellobiose were then incubated with the resin, in order to allow proteins able to interact with ClSca to bind to it while bound to the resin. The resin was then washed and ClSca, and any interacting partners were eluted. The eluted samples were concentrated ∼10-fold and analysed by SDS-PAGE.

These proteins are shown in Supplementary Table 3. After intuitive curation of this list, removing homologs of commonly identified housekeeping proteins (*e*.*g*. EF-Tu), two proteins were examined further. These two proteins are encoded on the putative genes Clole_0709 and Clole_2778. Using a conserved domain BLAST search, it was revealed that Clole_0709 encodes a putative cellulase with an enzymatic domain and a bacterial immunoglobulin domain type VII (BIg-7 domain). Using AlphaFold^21,22^ (accessed via ColabFold^23^), the structure of this BIg-7 domain was predicted. Testing the hypothesis that this structure may echo that of a characterised dockerin, the output was then used in a Dali structural search^24^. The closest match was that of fibronectin(III) domains from CbhA of *A*.*thermocellus* (PDB: 3PDD) which have been show to aid the hydrolysis of cellulose by modifying its surface^25^.

This bioinformatic process was also conducted with the sequence of that of Clole_2778, which showed no identifiable putative domains via conserved domain BLAST search, however the AlphaFold predicted structure prediction is visually reminiscent of that of a dockerin. A structural alignment of this with the most similar structure identified by Dali is shown in Figure 6B (8% identity, Epstein-Barr Virus Nuclear Antigen 1, PDB 5WMF), and with a canonical dockerin domain from *A. thermocellus* (PDB: 1OHZ) in Figure 6C.

**Figure 6.**
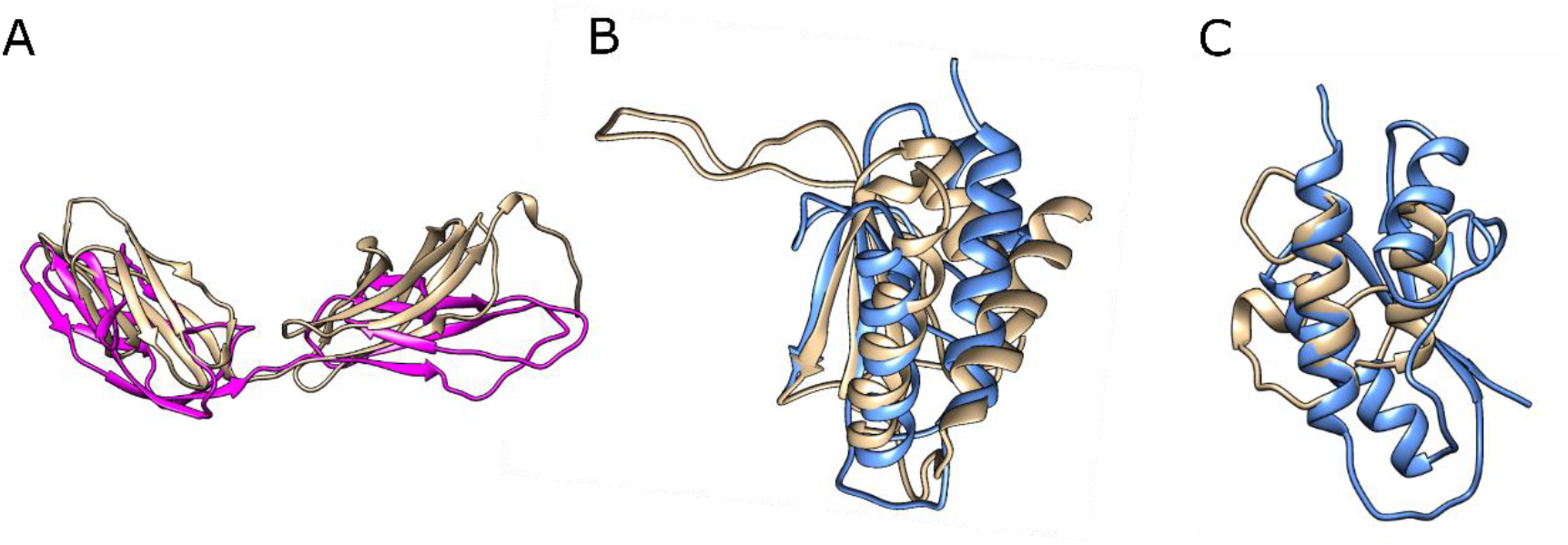
Novel interacting partners for ClSca align with existing crystal structures. The sequences of hits Clole_0709 (pink) and Clole_2778 (blue) were used as queries for AlphaFold structure prediction. The output structures were then used as queries in a DALi PDB search. Clole_0709 (pink) was most closely matched with the fibronectin-like domains of CbhA in structure 3PDD, and Clole_2778 (blue) most closely with Epstein-Barr Nuclear Antigen-1 (5WMF). Alignments with these structures are shown in respectively in A and B produced with UCSF Chimaera. The predicted structure of Clole_2778 was also used in a Chimaera MatchMaker alignment with the dockerin domain in structure 1OHZ (C).

## Discussion

Cellulosomes are normally enormous branching protein complexes that may include more than 130 enzyme subunits^13^. We have demonstrated that in the genome of *C. lentocellum*, a functional comparatively miniature cellulosome complex is encoded. Interestingly, recently, a small cellulosome was also discovered in *Ruminoclostridium saccharoperbutylacetonicum*^26^. The genome of this organism encodes two cohesin domains and eight dockerin-bearing enzymes. To our knowledge, the *C. lentocellum* cellulosome is the smallest currently described.

The closest neighbour of the cohesin domain from *C. lentocellum* is from *Cellulosilyticum* WCF-2. This domain is 100% identical (supplementary figure 1), however there is no biochemical characterisation of the domain. The genome of WCF-2 does not appear to encode any additional cohesin domains either. The lack of dockerin homologs in *Cellulosilyticum* WCF-2 presents the possibility that WCF-2 cellulosomes could cross-react and collaborate with *C. lentocellum* cellulosomes. There are no identifiable crystal structures which are homologous to the *C. lentocellum* cohesin domain when searching the PDB sequence database using BLAST, which presents the necessity for finding the structure of ClSca. Unfortunately, it was not possible to produce a crystal sufficient for X-ray crystallography (data not shown).

The dockerin domain of ClGH is >60% homologous to the dockerin domain of the canonical dockerin of *Acetivibrio thermocellus* (previously *Clostridium thermocellum*), with another close uncharacterised neighbour in *Ruminoclostridium cellobioparum*. The glutamate residues implicated in the cohesin dockerin interaction are conserved, so future studies should investigate the potential for the ClGH dockerin to interact with divergent cohesins.

The defining characteristic of cellulosomes is the interaction between cohesin and dockerin domains. The *C. lentocellum* system here encodes only one each of these on a discrete operon. Their inclusion on this operon, provided a suitable hypothesis that they would function together. By size exclusion chromatography we were able to purify a heterodimer of ClGH and ClSca. Assays of cellulase activity in the fractions obtained from this analysis also indicated the presence of ClGH in all collected fractions, including those under “Peak B” (Figure 5). The protein samples corresponding to this peak showed a protein of size smaller than ClGH by SDS-PAGE and larger than ClSca. Together with the evidence of cellulase activity, it is surmisable that this protein might be a cleaved version of ClGH, lacking it’s dockerin domain. There was no detectable ClSca un-dimerised however. Isothermal titration calorimetry was pursued, and titrations indicated an high affinity interaction, however optimal conditions were not achieved to permit calculation of biophysical constants (data not shown).

Classical enzyme kinetics experiments indicated that ClGH has relatively poor affinity for cellulose. The addition of ClSca however resulted in the formation of a complex with an enhanced specific activity (Figure 5). While this phenomenon is demonstrated consistently in other cellulosomes, it may also go some way to explaining why the organism has conserved such a small cellulosome. The CBM domain of the scaffoldin, which we have demonstrated in Figure 3 interacts with cellulose, is required for increased rates of cellulose breakdown.

The discovery here of additional interacting partners for ClSca may also be implicated in this regulation. Clole_0709 is predicted by dbcan to be a GH9 class enzyme. Other GH9 enzymes from cellulosomal organisms are endoglucanases (with 59% identity with an *A. thermocellus* GH9 endoglucanase), however other members of this class can also be cellobiohydrolases. Interestingly, the closest genetic relative of Clole_0709 also exists in WCF-2. The growing overlap in homology and repertoire of genes between these organisms presents further questions for biochemical study of a potentially divergent branch of cellulosomes, and interaction modes.

Other than the putative GH9 cellulase domain, Clole_0709 encodes a putative bacterial immunoglobulin group 7 (BIg-7) domain. This domain should be the focus of further study into the interaction of Clole_0709 and ClSca. Previous investigators have identified this domain in cellulosomal enzymes and solved structures of them^27^. These results reveal that the domain has some simplistic beta strands with coordinated Ca^2+^ ions. The structure does not appear similar at all to dockerin domains, however the coordination of these Ca^2+^ ions is shared. Interaction with cohesins was noted but has been left unexplored.

Besides Clole_0709, Clole_2778 should be studied further in its interaction with ClSca and other cohesins. It is not possible to predict any domains in the sequence of this small protein via conserved domain BLAST search, however AlphaFold predicts a structure with two antiparallel alpha helices which echo a dockerin domain. While Dali structural alignment search with PDB returned poorly aligned (8% at best) hits with proteins of eukaryotic viruses (Figure 6B), a structural alignment with dockerin domains is not unacceptable (Figure 6C).

The circumstances under which *C. lentocellum* produces this novel cellulosome should be investigated in future. Other cellulosomal organisms regulate cellulosome expression in response to cellobiose sensing at a cytoplasmic level^28^, and cell-surface mediated sensing of cellulose fibres^29,30^. These mechanisms exist however in organisms which have far more putative dockerin-bearing enzymes, of diverse functional classes. They are at least very different, if not also more complex. Cellobiose would be a natural instigator of cellulosome expression, given ease of passage to the cytoplasm, however the disparity in the apparent complexity of the cellulosomal system at this stage might indicate a different type of regulation. At present it may be sensible to surmise that the production of ClSca and ClGH is co-regulated with a variety of other non-cellulosomal genes, rather than with their own specific regulon. The discovery here though of putative additional members of the *C. lentocellum* cellulosome may begin to uncover more complexity in this regulation.

The current work provides useful biochemical characterisation of the *C. lentocellum* cellulosome. While this is the smallest cellulosome described to date, the identification of a novel enzyme which interacts with the *C. lentocellum* also presents a new line of investigation into other domains which might also perform this activity. Deeper biochemical investigation of Clole_0709 and Clole_2778 will describe the nature of their interaction with cohesin domains and compare this to conventional dockerins. This could indicate a hitherto hidden level of complexity in cellulosome structure and assembly. Crystal structures of the ClSca interacting with all three of these partners we have identified would be instrumental in uncovering this.

### Experimental Procedures

#### Gene discovery and cloning

Genes *clSca and clgh* were identified by PSI-BLAST searches of a pre-defined library of strains of the DSMZ GEBA collection of microorganisms. We hold a copy of this collection in our strain stocks. Query sequences used were alignments of the PFAM cohesin domain (PFAM_10345) group. This resulted in hits of non-novel domains from organisms in the collection such as *Acetivibrio thermocellum* (previously *Hungateiclostridium thermocellum*, and even more previously *Clostridium thermocellum*).

*C. lentocellum s*equences identified in this search were analysed using CLUSTALW to compare them to the canonical sequences present in PFAM for both cohesin and dockerin domains.

*C. lentocellum* was cultured for in peptone cellulose solution B (PCSb, modified from Lú *et al*_31_, 2 g/L peptone, 0.882g/L CaCl_2_.2H_2_O, 1 g/L yeast extract, 5 g/L NaCl) at 30 _o_C under anaerobic conditions with shaking at 200 rpm. Genomic DNA was prepared from the resulting culture using the Sigma GenElute Microbial DNA Extraction kit. Primers were generated which would amplify and add NdeI and HinDIII sites to the 5’ and 3’ ends of both ClGH and ClSca. After amplification of the genes with these primers from genomic DNA, the fragments were digested with the appropriate NEB branded enzymes and ligated using T4 DNA ligase (NEB) to the appropriate vector. After unsuccessful attempts to purify ClGH protein, a second plasmid was produced by deletion of the putative Sec signal peptide from the gene using the NEB Q5 site directed mutagenesis kit and a new set of primers. All primers and plasmid sequences are available in the supplementary materials.

#### Protein production and purification

ClGH was produced in BL21(DE3) *E. coli* by transformation of the pETClGHdsec plasmid and culture in autoinduction medium (Formedium) overnight at 30 °C with shaking at 200 rpm. ClSca was also produced in BL21(DE3) *E. coli*, transformed with the pQEClSca plasmid. These were cultured at 20 °C for 48 hours at 200 rpm in autoinduction medium. Generally, 500 mL cultures were utilised, which were split in to 50 mL aliquots from which bacteria were harvested and stored as pellets at -20 °C until such a time as their lysis and purification of the protein was required.

To purify protein, harvested bacteria were resuspended in 10 mL CDI buffer (50 mM Tris pH 8, 50 mM NaCl, 12 mM CaCl_2_) per g of cells. The resulting suspension was sonicated using a Branson Digital Sonifier, and centrifuged at 25,000 G to remove insoluble material. The supernatant was used as cell-free extract and applied to 5 mL TALON resin (Takara) equilibrated with CDI buffer. The resin was washed with CDI buffer, followed by CDI buffer with 10 mM imidazole. The protein was eluted in CDI buffer with 100 mM imidazole. Proteins were concentrated and buffer exchanged with Millipore centrifugal filter columns before any experiments.

Purification of the ClSca-ClGH heterodimer was performed by cleaning these resulting proteins further using a 200 pg HiLoad 16/600 Superdex column (Cytiva) for SEC. The proteins were mixed together and concentrated, before again being run on the same column.

#### Avicel pull downs

Purifed ClSca was incubated with Avicel at a range of concentrations in CDI buffer for around three hours. The Avicel was then removed from the suspension by centrifugation and washed with CDI buffer. The pellets were then mixed with one tenth volume of Laemmli buffer and boiled for ten minutes. After a brief centrifugation, the supernatant fractions were run on polyacrylamide gels.

### Enzymological methods

Purified ClGH was diluted in CDI buffer to 40 µM and incubated with 0.05 % (m/v) of the designated substrate also prepared in CDI buffer. To estimate the enzyme’s Vmax, reactions were performed at a range of concentrations with PASC. Reducing sugar assays were performed by the dinitrosalicylate method^32^. Cellobiose released from these assays was calculated using data obtained from standards and experimental samples on a Dionex ICS-5000 HPAEC system with a CarboPac PA-10 column, according to Munoz *et al*^33^.

### Baited cohesin pull downs

Cell-free extract from BL21(DE3) *E. coli* expressing the pQEClSca plasmid as described above was bound to 5 mL TALON resin. The resin was then washed with CDI buffer. *C. lentocellum* was grown in 50 mL PCSb overnight at 30 °C and the cells harvested. Cell-free extract was prepared from these cells as described above. The CFE was then incubated with the resin containing the bound ClSca, and washed again with CDI buffer. The ClSca and any interacting partners were eluted using CDI with 100 mM imidazole. The resulting samples were concentrated using Millipore centrifugal filter columns.

## Acknowledgements

The authors would like to thank Prof David Bolam at Newcastle University for the use of his ITC apparatus. Additionally, Prozomix should be thanked for the use of their Prozamigo software for searching the database of genomes held in the DSMZ’s GEBA collection of microbial strains, of which *C. lentocellum* was also obtained. The Authors would like to thank Dr Jose Muñoz for training on HPAEC apparatus and Dr Paul James for help with and supply of molecular weight standards with size exclusion chromatography. Dr Ciarán Kelly should also be thanked for helpful discussion.

