## Supplementary Figures for "A miniature cellulosome with novel interaction modes"

# A

```
ClGH      1   KGDLNGDQSINSTDYVLLKRHLLGVSLLTDIALKSADVDEDGAVNEGDLL  50
              +GDLNGD+ +N+ D  LLK HLLG  LLT  ALK+AD+D  G+V+  DLL
AtXynB  719   RGDLNGDKEVNALDLTLLKMHLLGSQLLTGDALKAADIDSSGSVDALDLL  768

ClGH     51   KLESYLL  57
              +L SYLL
AtXynB  769   RLRSYLL  775
```

# B

```
ClGH      2   GDLNGDQSINSTDYVLLKRHLLGVSLLTDIALKSADVDEDGAVNEGDLLK  51
              GD+NGD +INSTD  +LKR +L    LTD A    ADVD++G++N  D+L
RcGH9     5   GDVNGDGTINSTDLTMLKRSVLRAITLTDDAKARADVDKNGSINAADVLL  54

ClGH     52   LESYLL  57
              L  YLL
RcGH9    55   LSRYLL  60
```

# C

```
ClSca     1   FAMTLETLQLKAGEEAIIPIKLNNVSTGINNANLVFTYDKELIEVQEILA  50
              FAMTLETLQLKAGEEAIIPIKLNNVSTGINNANLVFTYDKELIEVQEILA
C. WCF-2 288  FAMTLETLQLKAGEEAIIPIKLNNVSTGINNANLVFTYDKELIEVQEILA  337

ClSca    51   GEIVPNSELSFKSAIHQTKGTFNLLFASAKQDGSDLITQSGDMVQVKIKA  100
              GEIVPNSELSFKSAIHQT+GTFNLLFASA+QDGSDLITQSGDMVQVKIKA
C. WCF-2 338  GEIVPNSELSFKSAIHQTEGTFNLLFASAEQDGSDLITQSGDMVQVKIKA  387

ClSca   101   KKDFTSAPFKVLSMKDIADAQLKKITVLFKVK  132
              KKDFTSAPFKVLSMKDIADAQLKKITVLFKVK
C. WCF-2 388  KKDFTSAPFKVLSMKDIADAQLKKITVLFKVK  419
```

# D

```
ClSca          1   FAMTLETLQLKAGEEAIIPIKLNNV-STGINNANLVFTYDKELIEVQEIL  49
                   F + +E      G+   IPIKL NV S GINN +    +D  ++EV ++
I. frigidare 518   FTIDIEIATASPGDTVTIPIKLINVPSAGINNCDFRVGFDSNVLEVVDVS  567

ClSca         50   AGEIVPNSELSFKSAIHQTKGTFNLLFASAKQDGSDLITQSGDMVQVKIK  99
                    G+I+P   L F S I+  +   + LFA  +Q G++LI   G
I. frigidare 568   TGDIIPRPALDFYSYINTEEEYVSFLFADEEQQGNNLIYDDG--------  609

ClSca        100   AKKDFTSAPFKVLSMKDIADAQLKKITV  127
                       FT+  F+VL+    A A L KITV
I. frigidare 609   ---VFTNIEFRVLAG---APAGLTKITV  631
```

**Supplementary Figure 1, *C. lentocellum* cohesin and dockerin domain alignments with close relatives.** BLASTp was utilised with the *C. lentocellum* cohesin and dockerin sequences as queries, searching the nr database. Alignments were then produced using Smith-Waterman method in Snapgene. (A) shows ClGH dockerin domain aligned with the *Acetivibrio thermocellus* XynB dockerin, and (B) with *Ruminoclostridium clariflavum* GH9 dockerin domain. In (C) ClSca's cohesin domain is aligned with that of *Cellulosilyticum sp. WCF-2*, and in (D) with that of *Iocasia Frigidaire*.

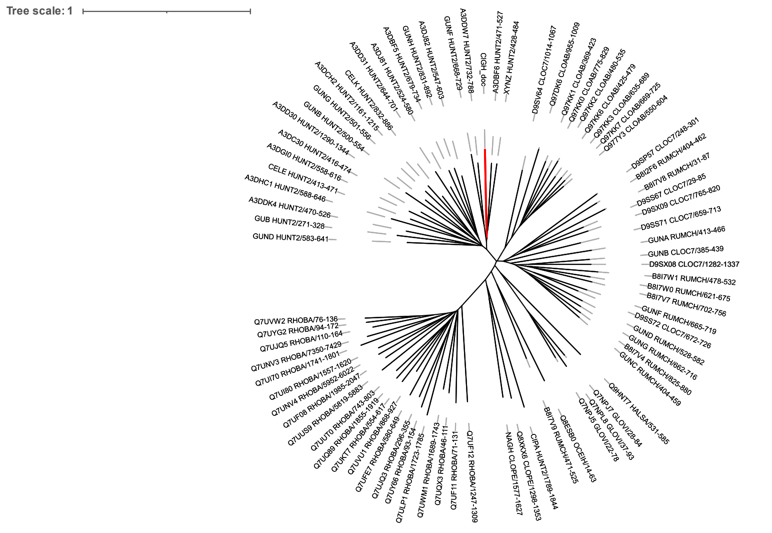


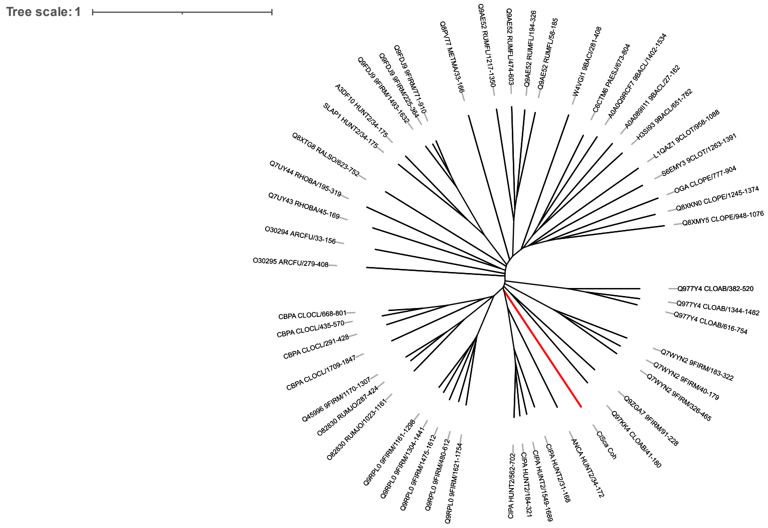


**Supplementary Figure 2 The *C. lentocellum* cohesin domain falls in to a distinct clade, while the dockerin is similar to those of *A. thermocellus*.** Utilising the sequences available in PFAM for dockerin (A) and cohesin (B) domains, phylogenetic trees were constructed. For dockerin domains, every single domain stored under this family in PFAM was utilised, however the diversity of cohesins was too large to perform this practically. The seed set of sequences was used instead. iTOL was used to producec unrooted phylogenetic trees with these sequences and the *C. lentocellum* genes are highlighted in bold red.
