## Supplementary Tables for "A miniature cellulosome with novel interaction modes"

| Protein names | Locus ID | Uniprot ID | Intensity |
|---|---|---|---|
| **ClSca** | **Clole_2599** | **F2JID6** | **1.28E+11** |
| Ribosomal RNA small subunit methyltransferase A (EC 2.1.1.182) | rsmA ksgA Clole_0654 | F2JN46 | 2.59E+09 |
| **Uncharacterized protein** | **Clole_2778** | **F2JK62** | **1.27E+09** |
| Elongation factor Tu (EF-Tu) | tuf Clole_3548 | F2JT13 | 3.88E+08 |
| Cadmium-translocating P-type ATPase (EC 3.6.3.3) | Clole_3672 | F2JH15 | 2.69E+08 |
| 50S ribosomal protein L6 | rplF Clole_3531 | F2JSH7 | 2.03E+08 |
| **Cellulase (EC 3.2.1.4)** | **Clole_0709** | **F2JNU4** | **1.79E+08** |
| Integral membrane sensor signal transduction histidine kinase | Clole_3138 | F2JP94 | 5.36E+07 |
| Histidine kinase (EC 2.7.13.3) | Clole_3860 | F2JJE5 | 2.62E+07 |
| Transcriptional repressor, CopY family | Clole_4056 | F2JL89 | 1.80E+07 |
| Transcription termination factor Rho (EC 3.6.4.-) (ATP-dependent helicase Rho) | rho Clole_1626 | F2JKU8 | 1.41E+07 |
| 60 kDa chaperonin (GroEL protein) (Protein Cpn60) | groL groEL Clole_3332 | F2JR41 | 1.39E+07 |
| Phosphoglycerate kinase (EC 2.7.2.3) | pgk Clole_2834 | F2JKZ2 | 1.23E+07 |
| ATP-dependent transcriptional regulator, MalT-like, LuxR family | Clole_3186 | F2JPU5 | 1.20E+07 |
| Signal recognition particle protein (Fifty-four homolog) | ffh Clole_2306 | F2JS77 | 1.13E+07 |

***Supplementary Table 3, Proteins identified in baited pull down experiment.*** From the baited pull-down experiment described in the results section, proteins obtained were subject to in-gel digestion with trypsin and then analysed by mass spectrometry. Peptides were annotated using MaxQuant with the Uniprot predicted proteome for this organism (UP000008467). Hits with intensities of less than $10^6$ were discarded. The remaining hits are listed in this table, in order of intensity. Those of note are displayed in bold.
